# Phylogenetic analysis of the origin and spread of plague in Madagascar

**DOI:** 10.1101/2022.03.29.486186

**Authors:** Luis R. Esquivel Gomez, Cyril Savin, Voahangy Andrianaivoarimanana, Soloandry Rahajandraibe, Lovasoa Nomena Randriantseheno, Zhemin Zhou, Arthur Kocher, Minoarisoa Rajerison, Xavier Didelot, Denise Kühnert

## Abstract

**Background:** Plague is a zoonotic disease caused by the bacterium *Yersinia pestis*, highly prevalent in the Central Highlands, a mountainous region in the center of Madagascar. After a plague-free period of over 60 years in the northwestern coast city of Mahajanga, the disease reappeared in 1991 and caused several outbreaks until 1999. Previous research indicates that the disease was reintroduced to the city of Mahajanga from the Central Highlands instead of reemerging from a local reservoir. However, it is not clear how many reintroductions occurred and when they took place.

**Methodology/Principal findings:** In this study we applied a Bayesian phylogeographic model to detect and date migrations of *Y. pestis* between the two locations that could be linked to the re-emergence of plague in Mahajanga. Genome sequences of 300 *Y. pestis* strains sampled between 1964 and 2012 were analyzed. Four migrations from the Central Highlands to Mahajanga were detected. Two resulted in persistent transmission in humans, one was responsible for most of the human cases recorded between 1995 and 1999, while the other produced plague cases in 1991 and 1992. We dated the emergence of the *Y. pestis* sub-branch 1.ORI3, which is only present in Madagascar and Turkey, to the beginning of the 20^th^ century, using a Bayesian molecular dating analysis. The split between 1.ORI3 and its ancestor lineage 1.ORI2 was dated to the second half of the 19^th^ century.

**Conclusions/Significance:** Our results indicate that two independent migrations from the Central Highlands caused the plague outbreaks in Mahajanga during the 1990s, with both introductions occurring during the early 1980s. They happened over a decade before the detection of human cases, thus the pathogen likely survived in wild reservoirs until the spillover to humans was possible. This study demonstrates the value of Bayesian phylogenetics in elucidating the re-emergence of infectious diseases.

**Author summary:** In 1991 human cases of plague were reported in the city of Mahajanga, located in the west coast of Madagascar, after 60 years without human cases. Existing evidence suggests that *Yersinia pestis*, the causal agent of the disease, was reintroduced to the city from a mountainous region known as the Central Highlands. We performed a phylogeographic analysis on 300 *Y. pestis* genome sequences to determine how many migrations of the pathogen between the two locations were related to the reappearance of the disease in Mahajanga. The results revealed that two migrations from the Central Highlands were the cause of the outbreaks of plague in the west coast of the country during the 1990s. We also aimed to date the emergence of the *Y*.*pestis* variant that circulates in Madagascar and is also present in Turkey. To do this, we conducted a molecular dating analysis using an extended data set of 445 sequences, which contained sequences from Turkey and India (the country from which the pathogen was exported to Madagascar for the first time). The analysis indicated that this particular variant emerged in the first decade of the 20^th^ century.

## Introduction

The gram-negative bacterium *Yersinia pestis* is the causal agent of plague, a zoonotic disease transmitted by hematophagous vectors (fleas). Despite being primarily a rodent disease, plague has infected humans for thousands of years [*1,2*], and has been responsible for at least three severe pandemics throughout human history: the first pandemic or Justinian plague (541-750 CE), the second pandemic that started with the Black Death (14^th^-18^th^ Century CE) and the third pandemic that originated in China in 1855 and spread globally from 1894 [*3*]. Altogether, plague claimed millions of lives during the course of these pandemics.

Currently, over 90% of the plague cases occur in the African continent [*4*]. The most affected country is Madagascar, an island located on the eastern coast of southern Africa, which accounts for 75% of all infections worldwide [*5*]. The strains of *Y. pestis* circulating in Madagascar belong to the 1.ORI3 phylogenetic sub-branch, which is part of the *Y. pestis* lineage responsible for the third plague pandemic [*3*]. According to historical records, plague was introduced in Madagascar when steamboats coming from India arrived at the port city of Toamasina in November 1898 [*6,7*]. Phylogenetic evidence suggests that plague was exported from Madagascar to the Anatolian peninsula (modern day Turkey) [*3*].

In decades following its introduction to Madagascar, plague spread through the country establishing three infection foci. Two are located in the Northern and Central Highlands, regions with an altitude of 800 meters above sea level, while the third one is found in the city of Mahajanga, located on the west coast of the country [*8,9*]. After reaching the capital city Antananarivo in 1921, *Y. pestis* became established in the Central Highlands [*10*]. In this region, human cases occur every year, most of them reported during the rainy season, from October to April [*8,11*]. This period coincides with a peak in the population levels of *Y. pestis* vectors (the flea species *Xenopsylla cheopis* and *Synopsyllus fonquerniei)* and with the lowest abundance of black rats (*Rattus rattus)*, the main host of *Y. pestis* in the country [*9,12*]. In Mahajanga, the first cases of plague were reported in 1902, with outbreaks occurring in 1907 and during the 1920s [*7*]. After 1928, no human cases were recorded in the city for over six decades until the disease reappeared in August 1991 (Fig 1) [*13*]. Plague continued to have a strong presence in the city for the rest of the 1990s and caused annual outbreaks until 1999, with the exception of the years of 1993 and 1994 [*14*]. The high transmission season in humans during these outbreaks ranged from July to November, coinciding with a high abundance of fleas on Asian Shrews (*Suncus murinus*), an important reservoir of *Y. pestis* in Mahajanga [*9,15,16*].

**Fig 1.**
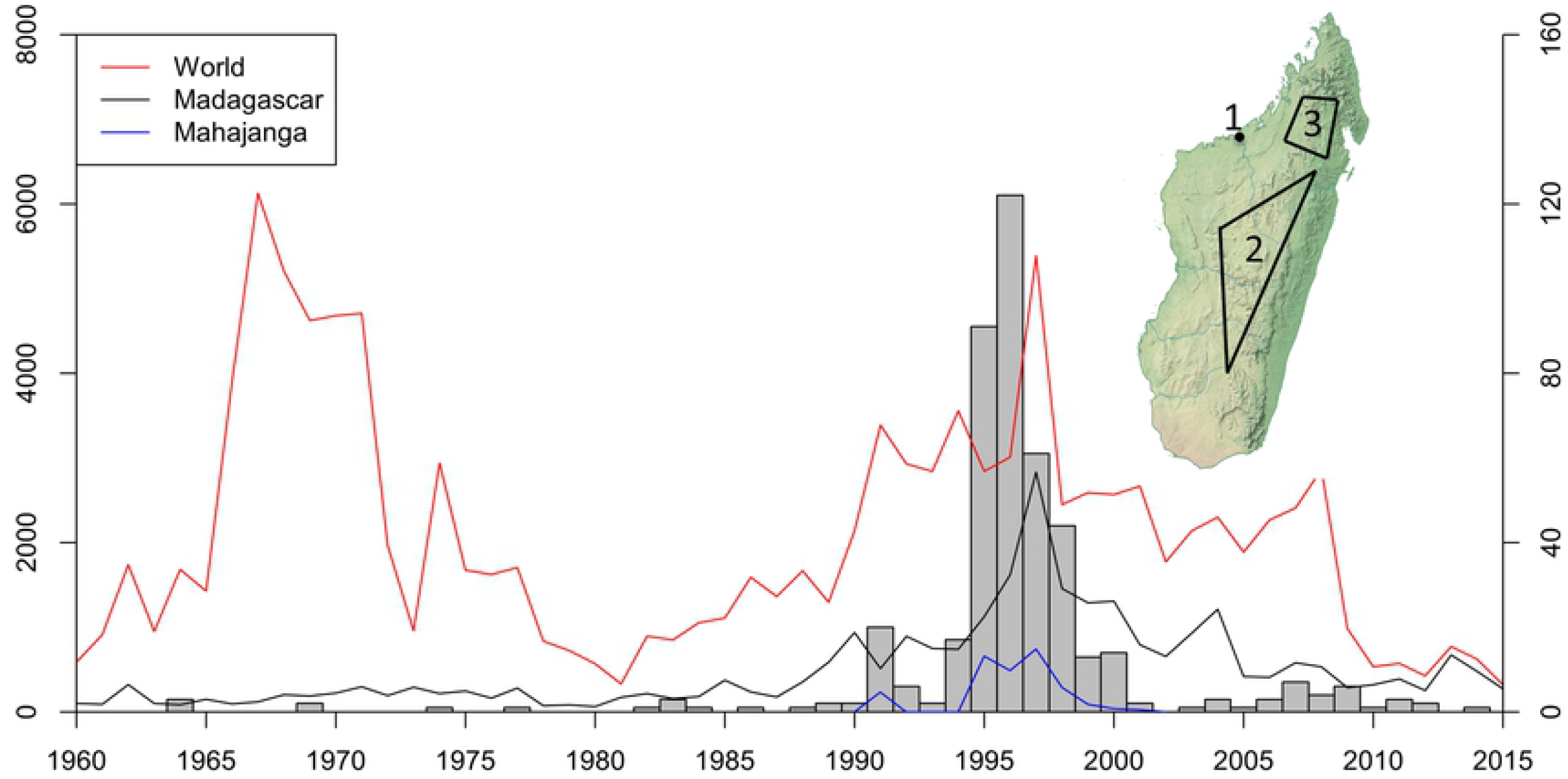
Reported plague cases worldwide, number of sequences used in the study and location of plague foci in Madagascar. Number of plague cases reported worldwide, for Madagascar and Mahajanga from 1960 to 2014 (left y-axis) [10]. The histogram represents the number of sequences used in the study (right y-axis). Upper right: Location of the three infection foci of plague in Madagascar. 1) Mahajanga, 2) Central Highlands, 3) Northern Highlands.

The sudden reappearance of plague in Mahajanga in 1991 raised the question of whether *Y. pestis* remained circulating in local wild reservoir populations for decades or if it was reintroduced to the city, presumably from the Central Highlands. Previous research based on the discovery of new single nucleotide polymorphisms (SNPs) found evidence of multiple transmissions of *Y. pestis* between the two locations, supporting the role of the Central Highlands as the source of the 1990s outbreaks [*9,17*]. While it is believed that a single genotype was responsible for the outbreaks in Mahajanga [*9*], it is not clear how many re-introductions of the bacterium were linked to the re-emergence of plague during the early 1990s, and how much time passed between the re-introduction of *Y. pestis* to Mahajanga and the detection of the first human case in 1991.

In this study, we aimed to better understand the migration dynamics of *Y. pestis* within Madagascar by performing a Bayesian phylogeographic analysis to estimate migration rates and the number and date of migration events of *Y. pestis* lineages between Mahajanga and the Central Highlands. We also performed a molecular dating analysis to estimate the time to the most recent common ancestor (tMRCA) of the 1.ORI3 strains present in Madagascar and Turkey, and to date the split between 1.ORI3 and its ancestral lineage 1.ORI2.a found in India. This allowed us to generate a timeline for the evolutionary epidemiology of *Y. pestis* starting with the split between 1.ORI2.a and 1.ORI3, followed by the introduction of the bacterium to India, Madagascar and subsequently to Turkey.

## Methods

### Data set

A total of 445 genomes of *Y. pestis* sampled between 1908 and 2014 from humans, rodents, fleas and shrews, were recovered from the online database EnteroBase (S1 Table) [*18*]. 440 genomes were sampled from Madagascar, three from Turkey and two from India. Sequences coming from samples that were cultured in laboratories for extended periods of time were not included in the data set as they likely present an excess of mutations. After the sequences were obtained, the EnteroBase Toolkit (EToKi) was used to align them against the *Y. pestis* reference genome CO92 (2001), to produce a SNP matrix. In brief, EToKi align module adopted minimap2 to align each genome against the CO92 genome, and extracted SNPs in the aligned regions. The same module also identified repeat regions including CRISPR regions, tandem repeats and interspersed repeats in the CO92 genome using PILER-CR, TRF and a self-comparison by minimap2, respectively. Any SNPs in the repeat regions, or in a region that was shared by ≤95% of genomes were discarded. The resulting core genomic, non-repetitive SNP matrix contained 783 sites (S2 Table).

While the genotype of the Indian sequences had been determined by a previous study [*3*], the presence in the matrix of 226 previously described SNPs related to 1.ORI3 (S3 Table) confirmed the genotype of the remaining sequences. This entire matrix was used to generate a time calibrated phylogeny of *Y. pestis*. For the phylogeographic analysis only sequences sampled from humans were used to avoid possible bias introduced by having samples from hosts with different travel ranges and capacities. This human-specific data set consisted of 300 sequences collected between 1964 and 2012, including 183 from the Central Highlands and 117 from Mahajanga.

### Test for phylogenetic signal

All Bayesian phylogenetic analyses were done with BEAST 2 [*19*], a platform developed for the inference of time calibrated trees. The evaluation of temporal signal for the two data sets was done with the BETS approach [*20*]. In this method, the marginal likelihoods of a heterochronous model (M1), with the real sampling dates, and an isochronous model (M2), with all dates set to zero and a fixed molecular clock rate, are estimated in a Path Sampling analysis [*21*]. A positive log Bayes Factor value (log marginal likelihood M1 - log marginal likelihood M2) indicates the presence of a temporal signal [*20,22*]. The Path Sampling analysis ran for 50 steps with a Markov Chain Monte Carlo (MCMC) length of 8,000,000 each.

### Phylogeographic analysis of *Y. pestis*

The phylogeographic analysis was done with the BEAST 2 package bdmm [*23*], which uses a structured (also called “multi-type”) birth–death model for the tree-generating process. Under this approach each sequence is assigned to a “type” or subpopulation and the type changes are estimated, following a type change rate *m*, jointly with the phylogenetic reconstruction. This analysis produces a multi-type tree, in which lineages of different types are annotated by different colors, and the migrations (type changes) are indicated by color changes occurring along the branches. Migration rates per lineage per year and the diversification-extinction ratio, which represents the number of new lineages generated per year and is also known as the reproductive number (R) in epidemiological contexts, are also estimated for each type.

We performed three replicate analyses with a MCMC of 300 million steps using a relaxed lognormal molecular clock and the HKY model of nucleotide substitutions. Sequences from Mahajanga were assigned to the type “0” and sequences from the Central Highlands to the type “1”. Given the evidence supporting the absence of plague in Mahajanga before 1991, we fixed the migration rates to 0 before 1980. This time point was selected since a significant increment in the number of plague cases in the Central Highlands started to occur during the early 1980s [*24*]. The same setup was applied for the estimation of the diversification-extinction ratio for Mahajanga (since there are no sequences sampled before 1991), however the value before 1980 was allowed to differ from 0 for the Central Highlands. In order to account for the uneven number of sequences per year observed in the data set, we estimated a different sampling proportion for each year and type. A Beta distribution with a mean equal to the number of samples/number of reported cases for each year, was chosen as the mean value of the Beta prior distribution used for each sampling proportion value. This allowed us to use plague incidence data to model the sampling process through time. Tip dates were sampled using a uniform distribution over the sampling year since it is very unlikely that all the sequences from one year were sampled on the same day. To improve MCMC mixing, the clock rate was fixed to a value of 2.85×10^−8^, obtained in a previous study for the branch 1 of *Y. pestis* [*25*]. As only SNP alignments were used, we corrected for ascertainment bias by specifying the number of invariable sites for each of the four bases, as found in the CO92 reference genome. The rate parameter for becoming uninfectious (δ) was fixed to a value of 1 to avoid identifiability issues with the other parameters [26].

The output files were combined with the software LogCombiner [*27*], after removing a 10% burn-in. This yielded effective sample sizes above the standard threshold of 200 for all parameters, as assessed by the software Tracer v.1.7 [*28*]. The maximum a posteriori tree (MAP), which represents the tree with the greatest posterior probability in the combined runs, was used for the visual representation of the migrations between foci, as it can display the type changes along a branch. The tree was visualized and edited with FigTree 1.4.4 [*29*]. As migration times on the MAP tree represent only point estimates, the median and 95% posterior intervals of migration times were calculated for each migration using a custom R script.

### Molecular dating analysis

The estimation of divergence times was done using a relaxed lognormal molecular clock, the HKY model of nucleotide substitutions and the Bayesian birth-death skyline tree prior implemented in the BEAST 2 package BDSKY [*26*]. The birth-death skyline model follows a forward in time birth-death process with only one lineage existing at time 0 in the past. As we move closer to the present, the lineages can split (birth event) at a rate λ, go extinct (death event) at a rate µ, and be sampled at a rate Ψ. These rates are reparametrized to estimate parameters like the sampling proportion (s), the become uninfectious rate (δ) and the diversification-extinction ratio [*26*]. The sampling proportion prior was specified per year, and tip date sampling and the correction for ascertainment bias were also applied in this analysis. The rate parameter for becoming uninfectious (δ) was also fixed to a value of 1.

Three replicate analyses were done with a MCMC length of 480 million states, and output files were later combined with LogCombiner, after removing a 10% burn-in. All parameters obtained effective sample sizes above 200. A maximum clade credibility tree was obtained for the combined runs using the program TreeAnnotator [*30*] with a 10% burn-in. The tree was edited with FigTree 1.4.4. The BEAST 2 XML files, output log files and the MAP and MCC tree files are provided as supporting information (S1-S6 Appendix).

## Results

### Migration dynamics of *Y. pestis*

We conducted a phylogeographic analysis based on the 300 human sequences from Madagascar. The temporal signal was significant for this subset with a BETS analysis giving a Bayes Factor of 52. According to the results of the bdmm analysis, the migration rate from the Central Highlands to Mahajanga after 1980 had a median value of 1.42×10^−3^ per lineage per year (95% HPD = 3.44×10^−4^ -3.05×10^−3^), and the rate from Mahajanga to the Central Highlands was 5.27×10^−4^ per lineage per year (95% HPD = 4.6×10^−6^ -2.04×10^−3^). The diversification-extinction ratio was significantly greater for Mahajanga with a median estimate of 5.2 (95% HPD = 4.68-5.76). For the Central Highlands the median values were 1.23 (95% HPD = 1.05-1.42) before 1980, and 2.96 (95% HPD = 2.65-3.26) afterwards (Table 1).

**Table 1.** Diversification-extinction ratio estimated for each infection focus, and migration rates of *Y. pestis* lineages per year from Mahajanga to the Central Highlands and vice-versa with the 95% highest posterior density intervals shown in brackets.

|  | **Diversification-extinction ratio** |
| --- | --- |
| Mahajanga | Post-1980: 5.2 [4.68- 5.76] |
| Central Highlands | Pre-1980: 1.23 [1.05-1.42]<br>Post-1980: 2.96 [2.65-3.26] |
|  | **Migration rates per lineage per year** |
| Mahajanga to Central Highlands | $5.27 \times 10^{-4}$ [$4.6 \times 10^{-6}$ - $2.04 \times 10^{-3}$] |
| Central Highlands to Mahajanga | $1.42 \times 10^{-3}$ [$3.44 \times 10^{-4}$ - $3.05 \times 10^{-3}$] |

The MAP tree used to summarize the migration history displayed five migration events, four of which were from the Central Highlands to Mahajanga (Fig 2). The first migration occurred around 1981 (95% HPD = 1980-1983) and originated a clade of sequences sampled in Mahajanga between 1995 and 1999. The second migration happened a year later (95% HPD = 1980-1984) and gave rise to a clade containing sequences sampled in 1991 and 1992. The additional migrations from the Central Highlands to Mahajanga had median estimates of 1989 (95% HPD = 1983-1997) and 1990 (95% HPD = 1985-1995) but each led to only a single sequence sampled in 1995 and 1997, respectively. Therefore, it seems that these introductions did not establish transmission chains in humans. The only migration from Mahajanga to the Central Highlands took place around 1996 (95% HPD = 1992-1997) leading to a single genome sampled in the capital Antananarivo in 1997.

**Fig 2.**
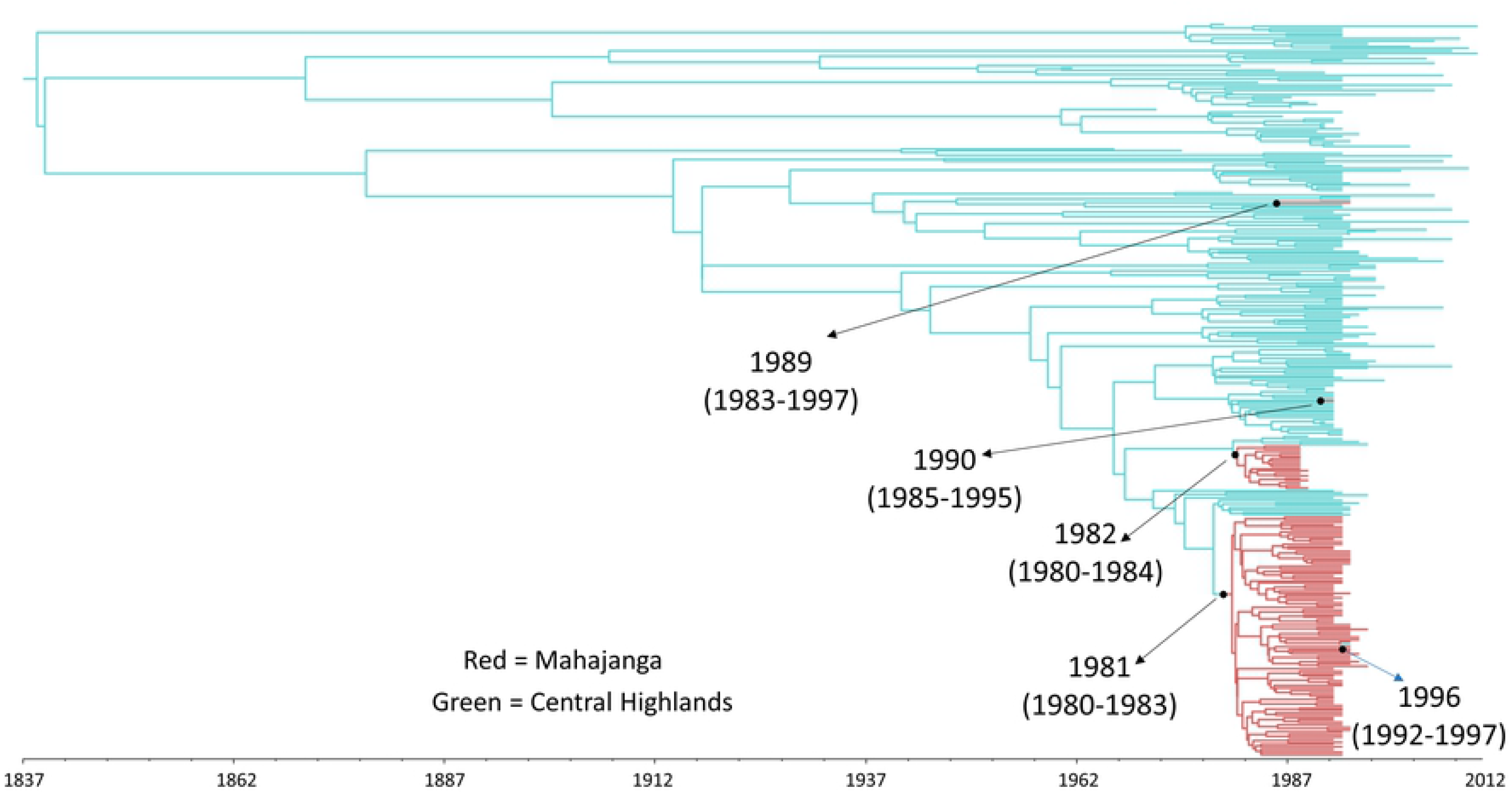
Maximum a posteriori typed tree of 300 *Y. pestis* sequences sampled in Mahajanga and the Central Highlands. Posterior median estimates for the dates of the migration events are shown with their respective 95% highest posterior density intervals (95% HPD). The four black arrows indicate migrations from the Central highlands to Mahajanga. The single blue arrow indicates a migration from Mahajanga to the Central Highlands.

### Molecular dating of *Y. pestis*

The BETS analysis for the full data set of 445 genomes of *Y. pestis* resulted in a log Bayes Factor of 203, which indicates a significant level of temporal signal. The BEAST 2 molecular dating analysis estimated a median evolutionary rate of 4.924×10^−8^ substitutions per site per year (95% HPD = 3.9×10^−8^ -6.11×10^−8^). The coefficient of variation of the clock rate had a median of 1.35 (95% HPD= 1.02-1.76), which indicates an important deviation from the strict clock model and a high degree of rate variation among the sequences. The tMRCA of all the sequences was estimated in 1877 (95% HPD = 1832-1904). The two Indian sequences shared a common ancestor in 1891 (95% HPD = 1859-1906). The most recent common ancestor of all sequences from Madagascar and Turkey (i.e. the 1.ORI3 sub-branch) was dated to 1905 (95% HPD = 1875-1927), while the divergence of the Turkish branch was dated to 1913 (95% HPD = 1887-1934). Lastly the three sequences from Turkey shared a most recent common ancestor in 1928 (95% HPD = 1906-1942) (Fig 3).

**Fig 3.**
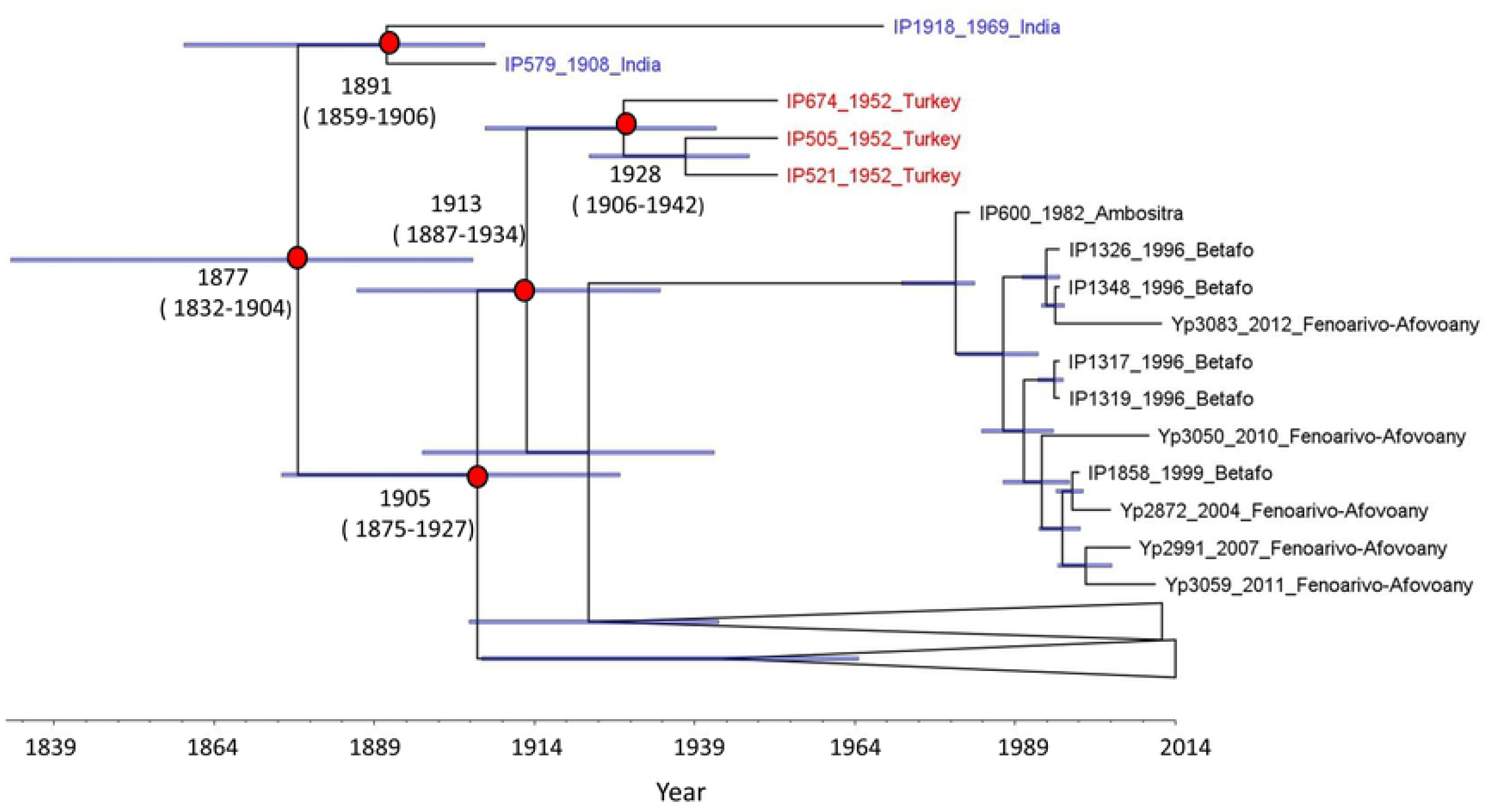
Maximum clade credibility tree of 445 sequences of *Yersinia pestis*. Sequences from India indicated with blue, sequences from Turkey with red. Posterior median estimates for the ages of the relevant nodes are shown with their respective 95% highest posterior density intervals (95% HPD). Error bars for the node heights are shown. Lower part of the phylogeny is collapsed to help with the visualization of the tree.

## Discussion

In this study we conducted a phylogeographic analysis of 300 *Y. pestis* genomes. The results revealed a higher migration rate of *Y. pestis* from the Highlands to Mahajanga, which supports the role of the former as the “source” of the *Y. pestis* populations that caused the outbreaks in the coastal city. Additional evidence in favor of a reintroduction rather than a reemergence of *Y. pestis* in Mahajanga include the lack of reports of massive deaths among rat populations during the period in which no human cases were reported in Mahajanga [*9*]. Also, while rodent populations of the black and brown rat (*Rattus norvegicus*) from the Central Highlands exhibit some degree of plague resistance, this is not the case for rats living in the west coast [*31*]. On the other hand, Asian shrews, currently the predominant plague reservoirs in Mahajanga exhibit a high resistance to *Y. pestis* [16].

Interestingly, most of the sequences from Mahajanga formed two distinctive clusters in the MAP tree, and each originated from different reintroductions of the pathogen that occurred in the early 1980s. This result suggests that two independent migrations were responsible for the establishment of the *Y. pestis* lineages that successfully started chains of transmission in humans. Curiously, it was the second migration event who originated the first outbreak of the decade, which started in August 1991 and ended seven months later [14], while the first migration of *Y. pestis* from the Central Highlands was the most successful, even if the introduced lineages jumped to human populations later. This result suggests that rodents infected by *Y. pestis* lineages introduced in the second migration were the first to reach, maybe just by chance, urban zones of Mahajanga that provided a good environment for rodent proliferation (e.g. abundant food, bad hygienic and economic conditions) and greater chances of close contact with humans. Both migrations occurred over a decade before the sampling of the oldest sequence in each clade, so it is very likely that the bacterium circulated in wild rodent or shrew populations before it jumped to humans, first in 1991 and a second time in 1995. After 1999, no human cases have been reported in Mahajanga, but unlike the disappearance of the disease after 1928, it is believed that *Y. pestis* continues to be present in the city, circulating in shrews with occasional spillover events to rats [*16*].

A previous study determined that there was a cycle of generation and loss of diversity in Mahajanga during the 1990s outbreaks with many lineages disappearing at the end of an epidemic year and new ones emerging during the next year [*9*]. This continued generation of lineages in Mahajanga could explain the greater diversification-extinction ratio found for this location compared to the Central Highlands. The greater diversification-extinction ratio of *Y. pestis* in Mahajanga, can also be related to the different environmental conditions of the two locations. The outbreaks in Mahajanga were centered around big markets with bad hygienic conditions, a good habitat to support a large number of rodents [*9*]. In the Highlands, plague is still considered a rural disease and the transmission of *Y. pestis* from rats to humans is linked in most cases to agricultural activities with low and high transmission seasons in humans present every year. The high transmission season in humans coincides with a peak in the flea population size and a low abundance of rats, as a result of the scarcity of food and the low temperatures that follow the harvest period, and the rat deaths caused by the disease itself [*13*]. Overall, it is believed that the maintenance of plague in Madagascar is the result of endemic cycles of transmission within the different districts of the country, mostly located in the Central Highlands [*32*].

While the results found here support previous claims of a reintroduction of *Y. pestis* to Mahajanga from the Central Highlands, it is also possible that additional migrations of this pathogen occurred from the Northern Highlands, the third plague focus in the country, and from other regions of Madagascar. In this regard, a recent study detected positive samples of *Y. pestis* in fleas sampled in the southwest part of the country [33]. While most of the samples were isolated from a flea species (*Echidnophaga gallinacea*) not suitable to be an efficient vector, the presence of the pathogen in an area far away from the plague foci suggests that the transmission of *Y. pestis* among non-human hosts is more widespread than previously thought. However, hypotheses of *Y. pestis* migrations to or from regions other than the Central Highlands couldn’t be tested with our dataset given the limited number of available sequences (<10) sampled outside the two main foci discussed here.

In this study we also generated a dated phylogeny of 445 *Y. pestis* sequences belonging to the ORI (Orientalis) biovar, collected from India (1.ORI2.a), Madagascar and Turkey (1.ORI3). The 1.ORI2.a sub-branch was exported from Hong Kong to India in 1896 [*3,34*] and represents the ancestor node of 1.ORI3, which is only found in Madagascar and Turkey. We estimated a median tMRCA for the 1.ORI2.a and 1.ORI3 sub-branches of 1877. This result indicates that the diversification of ORI likely occurred after the beginning of the third pandemic in 1855, as previously proposed [*3*], and was fueled by continuous exportations of the pathogen from Asia via sea trade.

The common ancestor of 1.ORI2.a was dated to 1891, 5 years before its arrival to India from Hong Kong. However, the 95% HPD intervals did contain the historical introduction year of 1896, supporting the arrival of plague to India at the end of the 19^th^ century. The estimate of 1905 for the most recent common ancestor of the sequences from Madagascar and Turkey suggests that the 1.ORI3 emerged around 7 years after the introduction of *Y. pestis* to the island in 1898. The appearance of this sub-branch within the first 10 years after the arrival of the pathogen to Madagascar could be the result of *Y. pestis* rapidly spreading and adapting to local reservoirs. The fact that no 1.ORI3 sequences have been detected outside Madagascar and Turkey, fit with a scenario in which 1.ORI2.a was introduced to Madagascar in 1898 with 1.ORI3 emerging within the next decade. A previous study estimated a median tMRCA of 1954 for Madagascan strains, although using only 33 genomes [*32*].

After reaching Madagascar, 1.ORI3 spread to Turkey probably in the early 1930s, as historical records mention the presence of two plague-infected individuals, coming from Madagascar, in the Middle East around 1931[*3*]. The results presented here support this claim, with the three sequences from Turkey having a median tMRCA of 1928. The presence of *Y. pestis* in samples from the 1950s, indicates that after its introduction, 1.ORI3 persisted (and likely continues to do so) in reservoir populations. The gerbils are potential candidates, as a few species have been recognized as important plague reservoirs in the neighboring country of Iran [*35*].

The evolutionary rate obtained in this study for the sub-branches 1.ORI2.a and 1.ORI3 is faster than the estimate for the Branch 1 of *Y. pestis* used in the phylogeographic analysis, which was obtained using sequences from the biovars 1.ANT (Antiqua), 1.IN (Intermediate) and a set of ancient genomes from the Second Pandemic [*25*], in addition to 1.ORI. This difference could be caused by the different time spans covered by the sequences in both studies. The sampling dates of the data set used in this study span 116 years, while some of the Second Pandemic genomes are over 600 years old. In this regard, a slower clock rate is expected in longer time spans as the effect of purifying selection becomes stronger [36]. This indicates that the slower rate, used to estimate the dates of the migrations, is likely closer to the true evolutionary rate of the branch 1 of *Y. pestis*. Moreover, assuming a higher evolutionary rate would not change the results of the phylogeographic analysis qualitatively.

Taking into account the results presented here, our study provides a timeline of the diversification of *Y. pestis* from the arrival of the 1.ORI2.a sub-branch to India in the late 19^th^ century to the spread of 1.ORI3 to Turkey a few decades later. This study, also presents the first application of a structured birth-death migration model to the study of *Y. pestis*, which allowed the dating of migration events and estimation of migration rates, while enabling the detailed modeling of the sampling process with the use of incidence data. Furthermore, if more *Y. pestis* sequences sampled from rodent and other small mammal species become available, a similar analysis could be done using a more complex model to account for the circulation of the bacteria in non-human hosts to assess their role in the spread and re-emergence of plague.

## Acknowledgments

We would like to thank Elisabeth Carniel for supervising the generation of the sequences, Mark Achtman for uploading of the sequences to Enterobase and the recovery of the sequences from the database. We thank Johannes Krause, Kirsten Bos, Alexander Herbig and Maria Spyrou for their valuable inputs and suggestions for the analyses, and Sanni Översti and Ariane Weber for their comments on the manuscript.

## Supporting information

**S1 Table. *Yersinia pestis* samples used in this study**

**S2 Table. SNP matrix generated with the 445 *Yersinia pestis* genomes**

**S3 Table. SNPs present in the 1.ORI3 subbranch of Yersinia pestis**

**S1 Appendix. BEAST2 xml file used for the bdmm analysis**

**S2 Appendix. Combined log file obtained with bdmm**

**S3 Appendix. MAP tree obtained with bdmm**

**S4 Appendix. BEAST2 xml file used for the BDSKY analysis**

**S5 Appendix. Combined log file obtained with BDSKY**

**S6 Appendix. MCC tree obtained with BDSKY**

